# Inhibition of Bruton’s tyrosine kinase activity attenuates trauma-induced multiple organ dysfunction in rats

**DOI:** 10.1101/2021.09.23.460775

**Authors:** Nikita M Patel, Filipe RMB Oliveira, Hanna Pillmann Ramos, Eleonora Aimaretti, Gustavo Ferreira Alves, Sina M Coldewey, Massimo Collino, Regina Sordi, Christoph Thiemermann

## Abstract

**Objective:** The aim of this study was to investigate (a) the potential of the Bruton’s tyrosine kinase (BTK) inhibitors (BTKi) acalabrutinib and fenebrutinib to reduce multiple organ dysfunction syndrome (MODS) in acute and chronic hemorrhagic shock (HS) rat models and (b) whether treatment with either acalabrutinib or fenebrutinib attenuates BTK, NF-κB and NLRP3 activation in HS.

**Background:** The MODS caused by an excessive systemic inflammatory response following trauma is associated with a high morbidity and mortality. The protein BTK is known to play a role in the activation of the NLRP3 inflammasome, which is a key component of the innate inflammatory response. However, its role in trauma-hemorrhage is unknown.

**Methods:** Acute and chronic HS rat models were performed to determine the influence of acalabrutinib or fenebrutinib on MODS. The activation of BTK, NF-κB and NLRP3 pathways were analyzed by western blot in the kidney.

**Results:** We demonstrated that (a) HS caused organ injury and/or dysfunction and hypotension (post resuscitation) in rats, while (b) treatment of HS-rats with either acalabrutinib or fenebrutinib attenuated the organ injury and dysfunction in acute and chronic HS models and (c) reduced the activation of BTK, NF-κB and NLRP3 pathways in the kidney.

**Conclusion:** Our results point to a role of BTK in the pathophysiology of organ injury and dysfunction caused by trauma/hemorrhage and indicate that BTK inhibitors may be repurposed as a potential therapeutic approach for MODS after trauma and/or hemorrhage.

**MINI-ABSTRACT:** This study evaluated the role of Bruton’s tyrosine kinase (BTK) in trauma/hemorrhage. Patients with trauma had elevated gene expression of BTK. The BTK inhibitors acalabrutinib (irreversible) and fenebrutinib (reversible) attenuated the trauma-induced multiple organ dysfunction in rats with hemorrhagic shock, indicating that BTK could be a potential therapeutic target.

## INTRODUCTION

Trauma is one of the leading causes of death and disability in those aged under 44 and exceeds the number of deaths caused by HIV, tuberculosis and malaria combined^1^. Globally, there are approximately 6 million trauma-related deaths each year. In contrast to most other diseases, instances of trauma and the accompanying mortality rates are rising. Trauma-associated hemorrhage accounts for almost 40% of all trauma mortalities and is frequently considered the biggest cause of preventable deaths in the world^2^. Trauma patients, who survive the initial injury, often develop multiple organ dysfunction (MODS) at a later stage^3^. Whilst the underlying mechanisms contributing to MODS have not been fully elucidated, it is thought that a) an excessive systemic inflammatory response secondary to the release of damage-associated molecular patterns (DAMPs) from extensive tissue damage and b) ischemia-reperfusion (I/R) injury play an important role^4^. At present, there are no specific pharmacological interventions used clinically to prevent the onset of MODS associated with HS.

Bruton’s tyrosine kinase (BTK) is a cytoplasmic, non-receptor protein tyrosine kinase belonging to the Tec family of kinases and was first discovered in X-linked agammaglobulinemia^5^. All cells of hematopoietic origin except plasma cells, natural killer cells and T-lymphocytes express BTK^6^. Whilst BTK was initially known for its critical role in B-lymphocyte development and, thus, adaptive immunity, more recent studies point to a pivotal role for BTK in innate immunity^7^.

It is known that trauma leads to a so-called ‘genomic storm’ that results in a change in >80% of cellular functions and pathways^8^. Specifically, there is an increased expression in genes related to systemic inflammatory and innate immune responses. Of particular interest was the observation that B-lymphocyte receptor signaling was one of the most upregulated pathways. We have recently discovered that inhibition of BTK activity with either ibrutinib or acalabrutinib reduces the multiple organ (cardiac, renal, hepatic) injury/dysfunction caused by cecum-ligation and puncture (CLP)-sepsis in the mouse. Moreover, the BTKi used reduced the activation of NF-κB and the NLRP3 inflammasome as well as the release of many pro-inflammatory cytokines and chemokines triggered by CLP-sepsis^9^. Most notably, X-Linked immunodeficient mice with inactive BTK function were also protected from sepsis-induced MODS^10^. Furthermore, prevention of NF-κB activation reduces the onset of multiple organ injury and improves the survival rate in rodent models of septic shock^11^, whilst pharmacological blockade of NF-κB activation using inhibitors of IκB kinase reduced the MODS caused by CLP^12^ and HS^13^.

Driven by the COVID-19 pandemic, there has been a significant focus on interventions (repurposing) that dampen the cytokine storm and pulmonary injury linked to severe acute respiratory syndrome coronavirus 2 (SARS-CoV-2). Most notably, there is a positive correlation between disease severity and BTK activity following SARS-CoV-2 infection^14–16^. Thus, BTKi appear to reduce systemic inflammation and ongoing clinical trials will assess the potential impact of this repurposing strategy on outcome in patients with COVID-19 (ClinicalTrials.gov Identifier: NCT04382586, NCT04665115, NCT04439006, NCT04375397, NCT04528667 and NCT04440007).

BTKi are commonly used in patients for the treatment of B-lymphocyte malignancies such as chronic lymphocytic leukemia and mantle cell lymphoma. However, they have also received approval by the Food and Drug Administration (FDA) for use in patients with marginal zone lymphoma, small lymphocytic lymphoma, Waldenström’s macroglobulinemia and chronic graft versus host disease^17^. Given the evident protective effects of BTKi administration in sepsis and COVID-19, both of which display an excessive systemic inflammatory response, we wished to explore the potential of repurposing BTKi in trauma-hemorrhage.

There is limited information about the role of BTK in trauma^18^. Using the gene expression data in whole blood leukocytes of trauma patients^8^, we reanalyzed the expression of BTK. Having found a significant increase in BTK expression in patients with trauma, we then used a reverse translational approach to investigate the effect of the commercially available, irreversible BTKi acalabrutinib and the novel, reversible BTKi fenebrutinib on the organ injury/dysfunction in two rat models of HS. Having shown that both BTKi largely abolished the organ injury/dysfunction associated with hemorrhagic shock, we then investigated whether or not modulation of the BTK activity may affect the activation of selective inflammatory pathways, which are known to be involved in trauma/hemorrhage, specifically both the NF-κB and the NLRP3 inflammasome cascades.

## METHODS

### BTK gene expression in human whole blood

Original data was obtained under Gene Expression Omnibus (GEO) accession GSE36809, published by Xiao and colleagues^8^. RNA was extracted from whole blood leukocytes of severe blunt trauma patients (n = 167) over the course of 28 days and healthy controls (n = 37) and hybridized onto an HU133 Plus 2.0 GeneChip (Affymetrix) according to the manufacturer’s recommendations. The dataset was reanalyzed for BTK gene expression in these two groups.

### Use of Experimental Animals -Ethical Statement

For the acute HS model, all animal procedures were approved by the Animal Welfare Ethics Review Board (AWERB) of Queen Mary University of London and by the Home Office (License number PC5F29685). For the chronic HS model, all animal procedures were approved by the Universidade Federal de Santa Catarina Institutional Committee for Animal Use in Research (License number 7396250219) and are in accordance with the Brazilian Government Guidelines for Animal Use in Research (CONCEA). All *in vivo* experiments are reported in accordance to ARRIVE guidelines.

### Experimental Design

Male Wistar rats (for acute model: Charles River Laboratories Ltd., UK; for chronic model: Universidade Federal de Santa Catarina, Brazil) weighing 250-350 g were kept under standard laboratory conditions (12 h light/dark cycle with the temperature maintained at 19–22 °C) and received a chow diet and water *ad libitum*. All animals were allowed to acclimatize to laboratory conditions for at least one week before undergoing any experimentation. Acalabrutinib (3 mg/kg; Insight Biotechnology, UK) and fenebrutinib (3 mg/kg; Insight Biotechnology, UK) were separately diluted in 5 % DMSO + 95 % Ringer’s Lactate (vehicle) and rats were treated (i.v. in acute and i.p. in chronic model) upon resuscitation.

### Acute Hemorrhagic Shock Model

The acute hemorrhagic shock model was performed as previously described^19–21^. Briefly, fifty-four rats were anesthetized with sodium thiopentone (120 mg/kg i.p. initially and 10 mg/kg i.v. for maintenance as needed) and randomized into six groups: Sham + vehicle (n = 9); Sham + acalabrutinib (3 mg/kg; n = 8), Sham + fenebrutinib (3 mg/kg; n = 8), HS + vehicle (n = 9); HS + acalabrutinib (3 mg/kg; n = 10); HS + fenebrutinib (3 mg/kg; n = 10). Blood was withdrawn to achieve a fall in mean arterial pressure (MAP) to 35 ± 5 mmHg, which was maintained for 90 min. At 90 min after initiation of hemorrhage (or when 25% of the shed blood had to be reinjected to sustain MAP, resuscitation was performed with the shed blood over a period of 5 min. An infusion of Ringer’s lactate (1.5 mL/kg/h) was started as fluid replacement 1 h after resuscitation and was maintained throughout the experiment for a total of 3 h. At 4 h post-resuscitation, blood was collected for the measurement of biomarkers of organ injury/dysfunction (MRC Harwell Institute, Oxfordshire, UK). Sham-operated rats were used as control and underwent identical surgical procedures, but without hemorrhage or resuscitation. Detailed description of the acute hemorrhagic shock model and sample collection can be found in the supplemental section (Supplemental Figure 1A).

### Chronic Hemorrhagic Shock Model

Thirty-eight rats were administered analgesia with tramadol (10 mg/kg i.p.) 15 min prior to anesthesia induction with ketamine and xylazine (100 mg/kg and 10 mg/kg i.m. respectively) and randomized into four groups: Sham + vehicle (n = 9); Sham + acalabrutinib (3 mg/kg; n = 10); HS + vehicle (n = 10); HS + acalabrutinib (3 mg/kg; n = 9). Blood was withdrawn to achieve a fall in MAP to 40 ± 2 mmHg, which was maintained for 90 min. At 90 min after initiation of hemorrhage (or when 25% of the shed blood had to be reinjected to sustain MAP), resuscitation was performed with the shed blood over a period of 5 mins plus 1.5 mL/kg Ringer’s lactate. At 24 h post-resuscitation, blood was collected for the measurement of organ injury parameters (Hospital Universitário Professor Polydoro Ernani de São Thiago, Brazil). Sham-operated rats were used as control and underwent identical surgical procedures, but without hemorrhage or resuscitation. Detailed description can be found in the supplemental section (Supplemental Figure 1B).

### Western Blot Analysis

Semi-quantitative immunoblot analysis was carried out in kidney tissue samples as previously described^22^. The following antibodies were used: rabbit anti-total BTK, rabbit anti-NF-κB, rabbit anti-total IKKα/β, rabbit anti-Ser^176/180^ IKKα/β, mouse anti-Ser^32/36^ IκBα, mouse anti-total IκBα, (from Cell Signaling), rabbit anti-Tyr^223^-BTK, rabbit anti-NLRP3 inflammasome (from Abcam) and mouse anti-caspase 1 (p20) (from Adipogen). Detailed description of the method can be found in the supplemental section.

### Quantification of myeloperoxidase activity

Lung and liver tissue samples from the chronic HS model were homogenized in liquid nitrogen with a pestle and mortar. The homogenate was then centrifuged at 13,000 x g at 4 °C for 10 min and the supernatant was assayed for myeloperoxidase (MPO) activity by measuring the H_2_O_2_-dependent oxidation of 3,3,5,5-tetramethylbenzidine (TMB). MPO activity was determined colorimetrically using an ultra-microplate reader (EL 808, BioTek Instruments, INC, USA) set to measure absorbance at 650 nm. Total protein content in the homogenate was estimated using the BCA assay (Thermo Fisher Scientific, Rockford, IL), according to the manufacturer’s instructions. MPO activity was expressed as optical density at 650 nm per mg of protein.

### Statistical Analysis

All data in text and figures are expressed as mean ± SEM of *n* observations, where *n* represents the number of animals/experiments/subjects studied. Measurements obtained from the patient groups and sham, control, acalabrutinib and fenebrutinib treated animal groups were analyzed by one-way ANOVA followed by a Bonferroni’s *post-hoc* test on GraphPad Prism 8.0 (GraphPad Software, Inc., La Jolla, CA, USA). The distribution of the data was verified by Shapiro-Wilk normality test, and the homogeneity of variances by Bartlett test. When necessary, values were transformed into logarithmic values to achieve normality and homogeneity of variances. Differences were considered to be statistically significant when p<0.05.

## RESULTS

### BTK gene expression is elevated in trauma patients

Xiao and colleagues^8^ isolated whole blood leukocytes, extracted the total cellular RNA and performed genome-wide expression analysis by array from severe blunt trauma patient blood samples collected within 12 h and on Days 1, 4, 7, 14, 21 and 28 post-injury. Genome-wide expression in white blood cells from trauma patients was compared against matched healthy controls. Dataset is available under Gene Expression Omnibus (GEO) accession GSE36809. We reanalyzed this dataset for BTK expression in these groups. When compared to healthy controls, BTK expression was significantly elevated at all time points except 12 h (p<0.05; Figure 1). An initial peak was noted at Day 1 followed by a gradual decrease, however, BTK expression remained elevated at Day 28.

**Figure 1:**
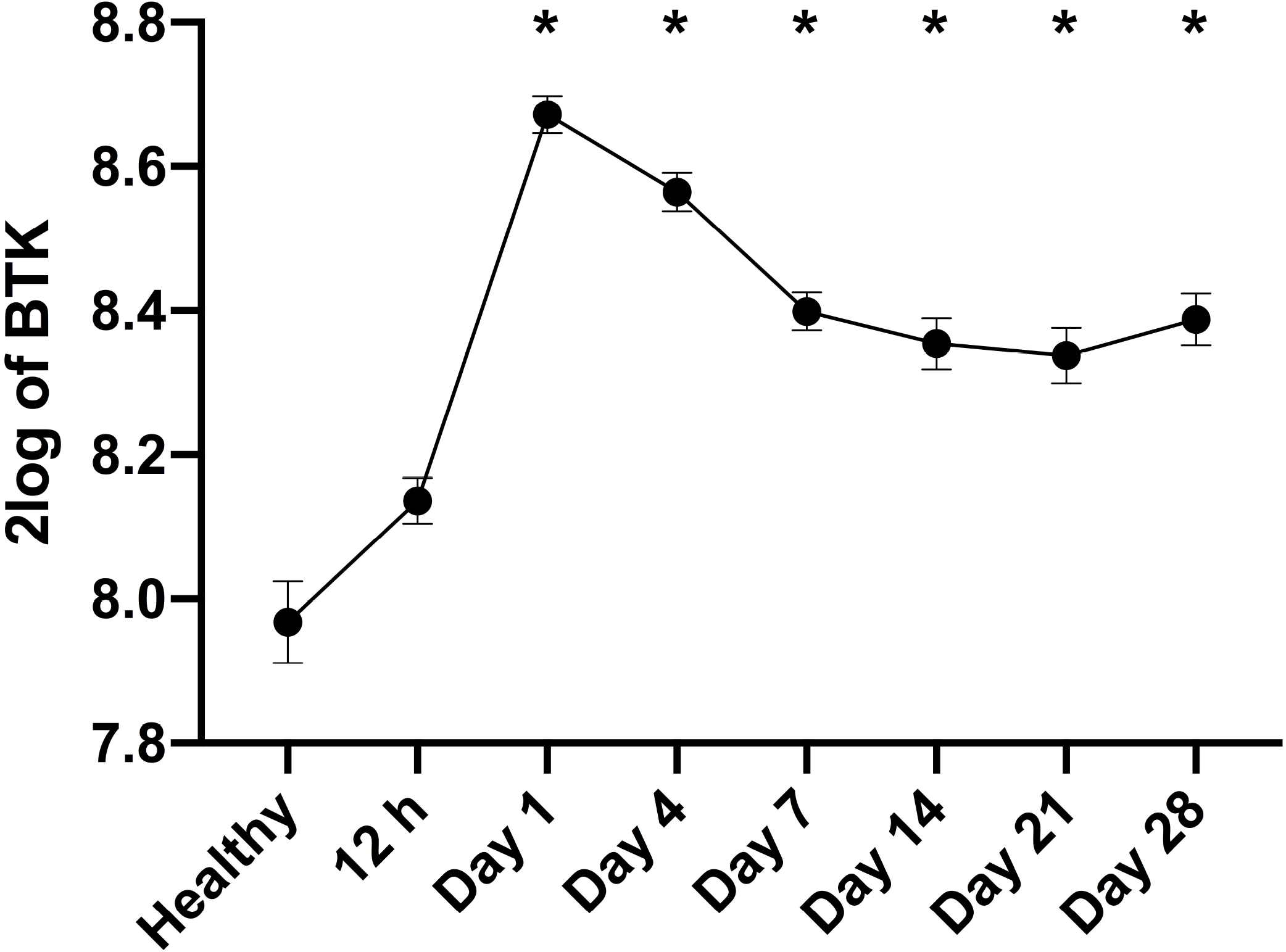
BTK gene expression is elevated in trauma patients. Original data was obtained from the Gene Expression Omnibus under dataset accession number GSE36809 which was published by Xiao and colleagues^8^. RNA was extracted from whole blood leukocytes over a 28-day time course from trauma patients (n = 167) and matched healthy controls (n = 37). Data were reanalyzed for BTK gene expression. Data are expressed as mean ± SEM. Statistical analysis was performed using one-way ANOVA followed by a Bonferroni’s *post-hoc* test. *p < 0.05 denoted statistical significance.

### Treatment with BTKi improves HS-induced circulatory failure in an acute HS model

To investigate the effects of the BTK inhibitors (BTKi) acalabrutinib and fenebrutinib on circulatory failure, MAP was measured from the completion of surgery to the termination of the experiment. Baseline MAP values were similar amongst all six groups. Rats subjected to HS demonstrated a decline in MAP which was ameliorated by resuscitation, but MAP remained lower than that of sham-operated rats during resuscitation (at the equivalent time points, Figure 2A). When compared to sham-operated rats, HS-rats treated with vehicle exhibited a more pronounced decrease in MAP over time post resuscitation. The MAP of HS-rats treated with either acalabrutinib or fenebrutinib was significantly higher than that of HS-rats treated with vehicle at the end of the resuscitation period (p<0.05; Figure 2B). No significant differences were observed between HS-rats treated with either acalabrutinib or fenebrutinib (p>0.05; Figure 2A). Administration of acalabrutinib or fenebrutinib to sham-operated rats had no significant effect on MAP (p>0.05; Figure 2A).

**Figure 2:**
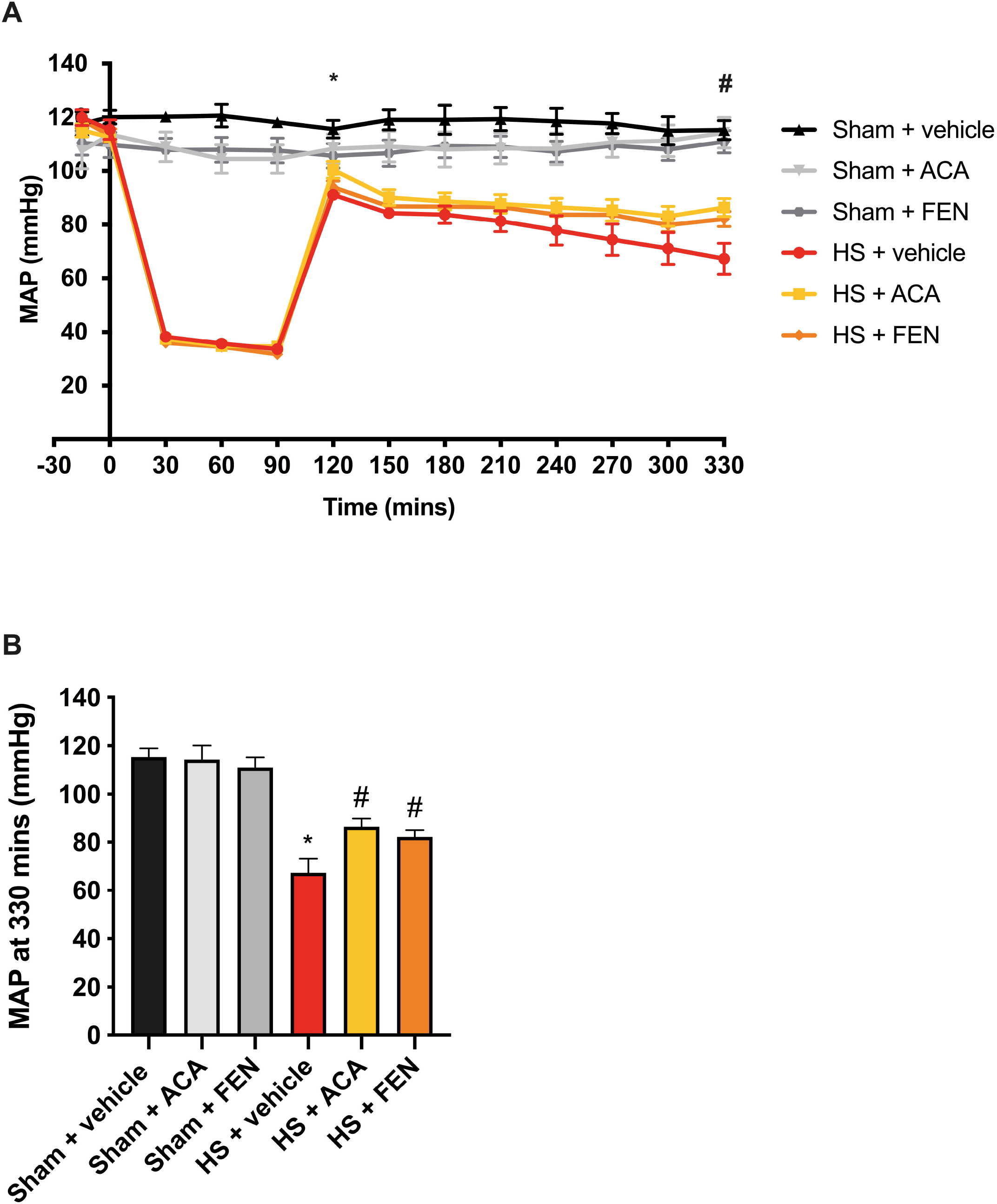
Treatment with BTKi improves HS-induced circulatory failure in an acute HS model. (**A**) Mean arterial pressure (MAP) was measured from the completion of surgery to the termination of the experiment for vehicle and BTKi treated (acalabrutinib, ACA; fenebrutinib, FEN) rats. (**B**) MAP values at the end of the resuscitation period (330 mins). Data are expressed as mean ± SEM of 8-10 animals per group. Statistical analysis was performed using two-way ANOVA followed by a Bonferroni’s *post-hoc* test. *p<0.05 Sham + vehicle vs. HS + vehicle; #p<0.05 HS + vehicle vs. HS + BTKi (ACA or FEN).

### Treatment with BTKi attenuates HS-induced organ damage in an acute HS model

Here we explored whether pharmacological intervention with the BTKi acalabrutinib and fenebrutinib attenuates the MODS associated with HS in rats. When compared to sham-operated rats, rats subjected to HS and treated with vehicle displayed increases in serum urea (p<0.05; Figure 3A) and creatinine (p<0.05; Figure 3B); indicating the development of renal dysfunction. When compared to sham-operated rats, vehicle treated HS-rats exhibited significant increases in ALT (p<0.05; Figure 3C) and AST (p<0.05; Figure 3D) indicating the development of hepatic injury, while the increases in CK (p<0.05; Figure 3E) and amylase (p<0.05; not shown) denote neuromuscular and pancreatic injury, respectively. The significant increase in LDH (p<0.05; Figure 3F) in HS-rats treated with vehicle confirms that tissue injury had occurred. Treatment of HS-rats with either acalabrutinib or fenebrutinib significantly attenuated the renal dysfunction, hepatic injury, neuromuscular injury and general tissue damage caused by HS as shown by the reduction in serum parameter values (all p<0.05; Figures 3A-F). Treatment with either BTKi had no significant effect on pancreatic injury (p>0.05; not shown). No significant differences were observed between HS-rats treated with either acalabrutinib or fenebrutinib (p>0.05; Figure 3). Administration of either BTKi to sham-operated rats had no significant effect on any of the parameters measured (p>0.05; Figure 3).

**Figure 3:**
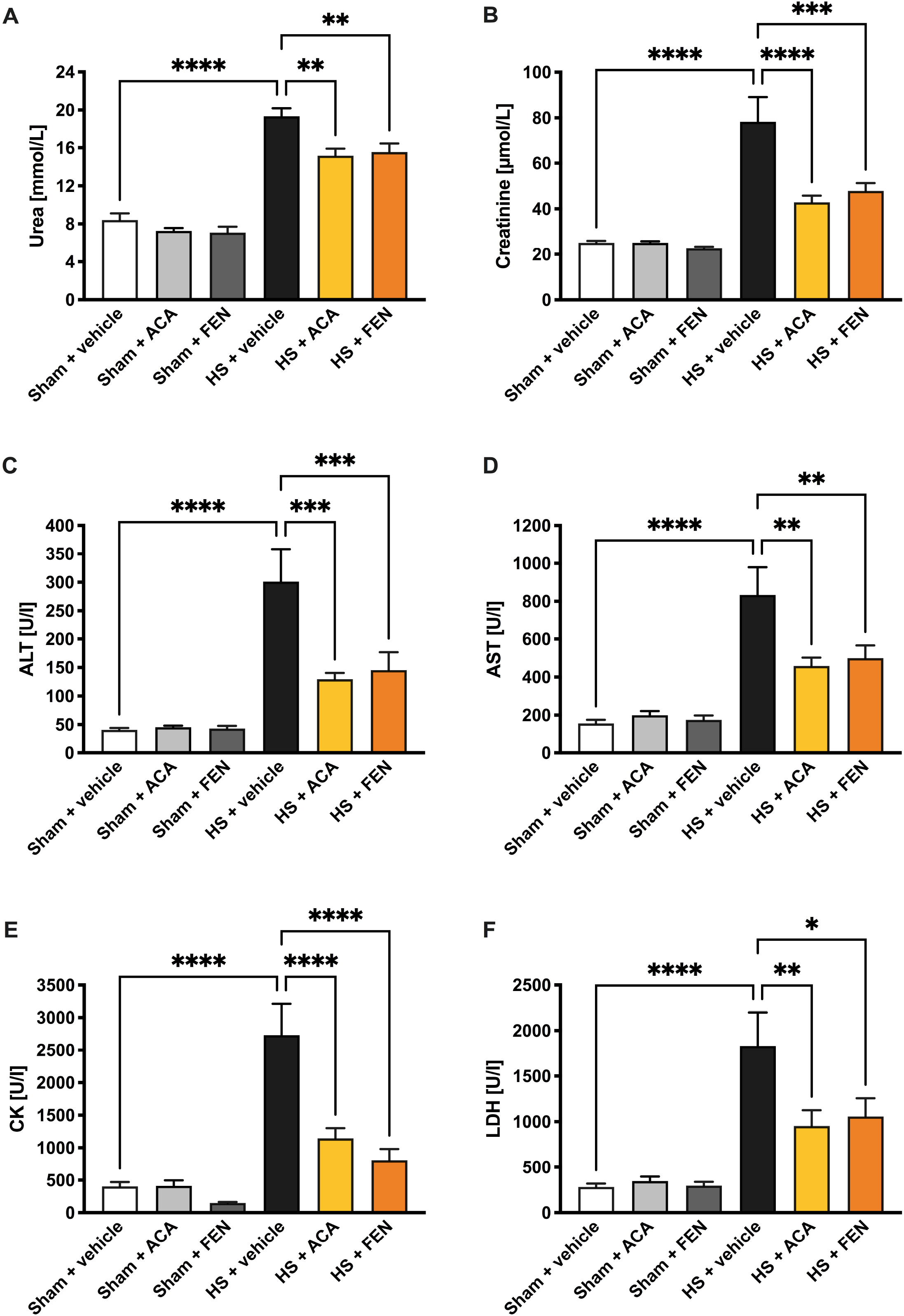
Treatment with BTKi attenuates HS-induced organ damage in an acute HS model. Rats were subjected to hemorrhagic shock (HS) and 4 h after resuscitation, levels of serum (**A**) urea, (**B**) creatinine, (**C**) alanine aminotransferase (ALT), (**D**) aspartate aminotransferase (AST), (**E**) creatine kinase (CK) and (**F**) lactate dehydrogenase (LDH) were determined in vehicle and BTKi treated (acalabrutinib, ACA; fenebrutinib, FEN) rats. Sham-operated rats were used as control. Data are expressed as mean ± SEM of 8-10 animals per group. Statistical analysis was performed using one-way ANOVA followed by a Bonferroni’s *post-hoc* test. *p<0.05 denoted statistical significance.

### Treatment with BTKi abolishes renal BTK activation in an acute HS model

Using western blot analysis, we examined whether HS leads to the activation of BTK in the kidney; given that treatment with either BTKi significantly attenuated HS-associated renal dysfunction. The activation of BTK and subsequent BTK-associated signaling pathways consists of the phosphorylation of BTK at Tyr^223^ as the initial stage of the BTK-signaling cascade. When compared to sham-operated rats, HS-rats treated with vehicle displayed significant increases in the phosphorylation of BTK at Tyr^223^, indicating that BTK is activated in injured kidneys (p<0.05; Figure 4A). Treatment with BTKi in HS-rats significantly abolished the increases in the phosphorylation of BTK at Tyr^223^ (p<0.05; Figure 4A). No significant differences were observed in the degree of phosphorylation (in the kidney) of BTK at Tyr^223^ between HS-rats treated with either acalabrutinib or fenebrutinib (p>0.05; Figure 4A). These data demonstrate that both BTKi abolish the activation of BTK caused by HS.

**Figure 4:**
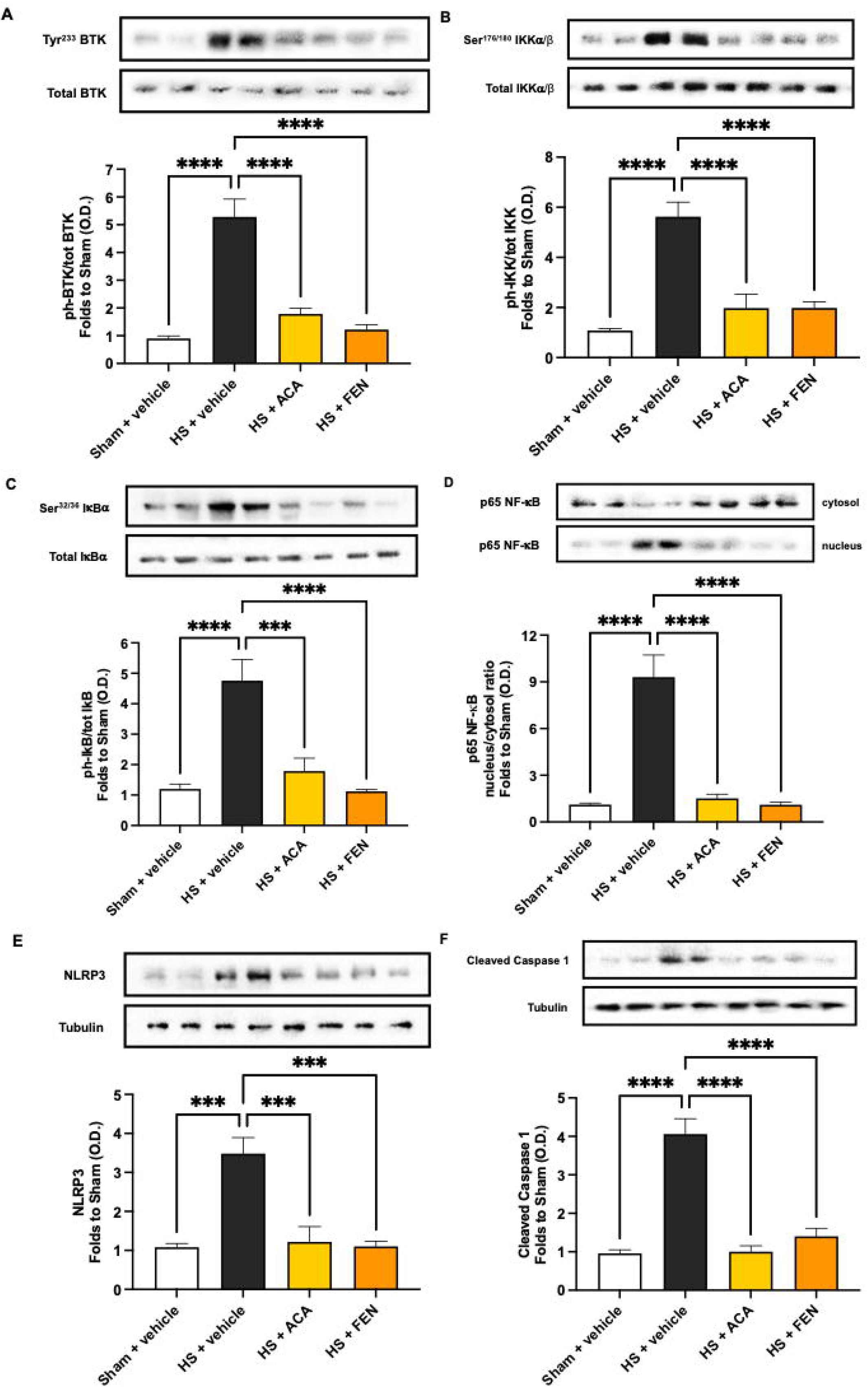
Treatment with BTKi attenuates BTK, NF-κB and NLRP3 activation in an acute HS model. (**A**) The phosphorylation of BTK at Tyr^223^, (**B**) the phosphorylation of IKKα/β at Ser^176/180^, (**C**) the phosphorylation of IκBα at Ser^32/36^, (**D**) the nuclear translocation of p65, (**E**) the activation of NLRP3 and (**F**) the cleaved (activated) form of caspase 1 of vehicle and BTKi treated (acalabrutinib, ACA; fenebrutinib, FEN) rats were determined by western blotting in the kidney. Protein expression was measured as relative optical density (O.D.) and normalized to the sham band. Data are expressed as mean ± SEM of five animals per group. Statistical analysis was performed using one-way ANOVA followed by a Bonferroni’s *post-hoc* test. *p<0.05 denoted statistical significance.

### Treatment with BTKi abolishes renal NF-κB activation in an acute HS model

The effect of BTK inhibition on the activation of the signaling events leading to the activation of NF-κB, were investigated in the kidney. When compared to sham-operated rats, HS-rats treated with vehicle had significant increases in the phosphorylation of IKKα/β at Ser^176/180^ (p<0.05; Figure 4B), phosphorylation of IκBα at Ser^32/36^ (p<0.05; Figure 4C) and the translocation of p65 to the nucleus (p<0.05; Figure 4D). Treatment of HS-rats with either BTKi significantly abolished the increase in renal phosphorylation of IKKα/β at Ser^176/180^ (p<0.05; Figure 4B), the phosphorylation of IκBα at Ser^32/36^ (p<0.05; Figure 4C) and the nuclear translocation of p65 (p<0.05; Figure 4D). No significant differences were observed in the degree of phosphorylation of IKKα/β at Ser^176/180^, phosphorylation of IκBα at Ser^32/36^ and the translocation of p65 to the nucleus between HS-rats treated with either acalabrutinib or fenebrutinib (p>0.05; Figures 4B-D). These data illustrate that both BTKi abolish the activation of NF-κB caused by HS.

### Treatment with BTKi abolishes renal NLRP3 and caspase 1 activation in an acute HS model

Having discovered that BTKi significantly reduced the activation of NF-κB in the kidney of rats subjected to HS, we next analyzed the potential involvement of the NLRP3 inflammasome complex. When compared to sham-operated rats, HS-rats treated with vehicle exhibited a significantly increased expression of the NLRP3 inflammasome (p<0.05; Figure 4E) and of the cleaved (activated) form of caspase 1 (p<0.05; Figure 4F). Treatment of HS-rats with either BTKi acalabrutinib or fenebrutinib significantly inhibited the renal expression of NLRP3 (p<0.05; Figure 4E) and cleaved form of caspase 1 (p<0.05; Figure 4F). No significant differences were observed in the degree of expression of the NLRP3 inflammasome and cleaved form of caspase 1 between HS-rats treated with either acalabrutinib or fenebrutinib (p>0.05; Figures 4E-F). These data demonstrate that both BTKi abolish the activation of the NLRP3 inflammasome and caspase 1 and the subsequent formation of IL-1β.

### BTK activation correlates with renal dysfunction, NF-κB and NLRP3 activation in an acute HS model

Firstly, to address the question whether the degree of activation of BTK correlates with changes in renal function, we correlated the degree of phosphorylation of BTK at Tyr^223^ with serum creatinine (Figure 5A), urine creatinine (Figure 5B) and serum urea (Figure 5C). We found a highly significant positive correlation between the degree of BTK activation and the increases in serum creatinine (Figure 5A) and urea (Figure 5C), suggesting that BTK activation drives or precedes the renal dysfunction associated with hemorrhagic shock. No significant correlation was observed between BTK activation and the decrease in urine creatinine (Figure 5B). Secondly, the potential relationship between the degree of activation of BTK and alterations in the activation of NF-κB was also addressed by correlating the degree of phosphorylation of BTK at Tyr^223^ with the phosphorylation of IKKα/β at Ser^176/180^ (Figure 5D), the phosphorylation of IκBα at Ser^32/36^ (Figure 5E) and the translocation of p65 (Figure 5F). We found a highly significant positive correlation between the degree of BTK activation and NF-κB activation when measured as IKKα/β phosphorylation at Ser^176/180^ (Figure 5D), IκBα phosphorylation at Ser^32/36^ (Figure 5E) and p65 translocation (Figure 5F). Thirdly, whether the degree of activation of BTK correlates with changes in the assembly and activation of the NLRP3 inflammasome was investigated by correlating the degree of phosphorylation of BTK at Tyr^223^ with the expression of NLRP3 (Figure 5G) and the cleaved (activated) form of caspase 1 (Figure 5H). We found a highly significant positive correlation between the degree of BTK activation and the NLRP3 inflammasome expression (Figure 5G) and the activation of caspase 1 (Figure 5H).

**Figure 5:**
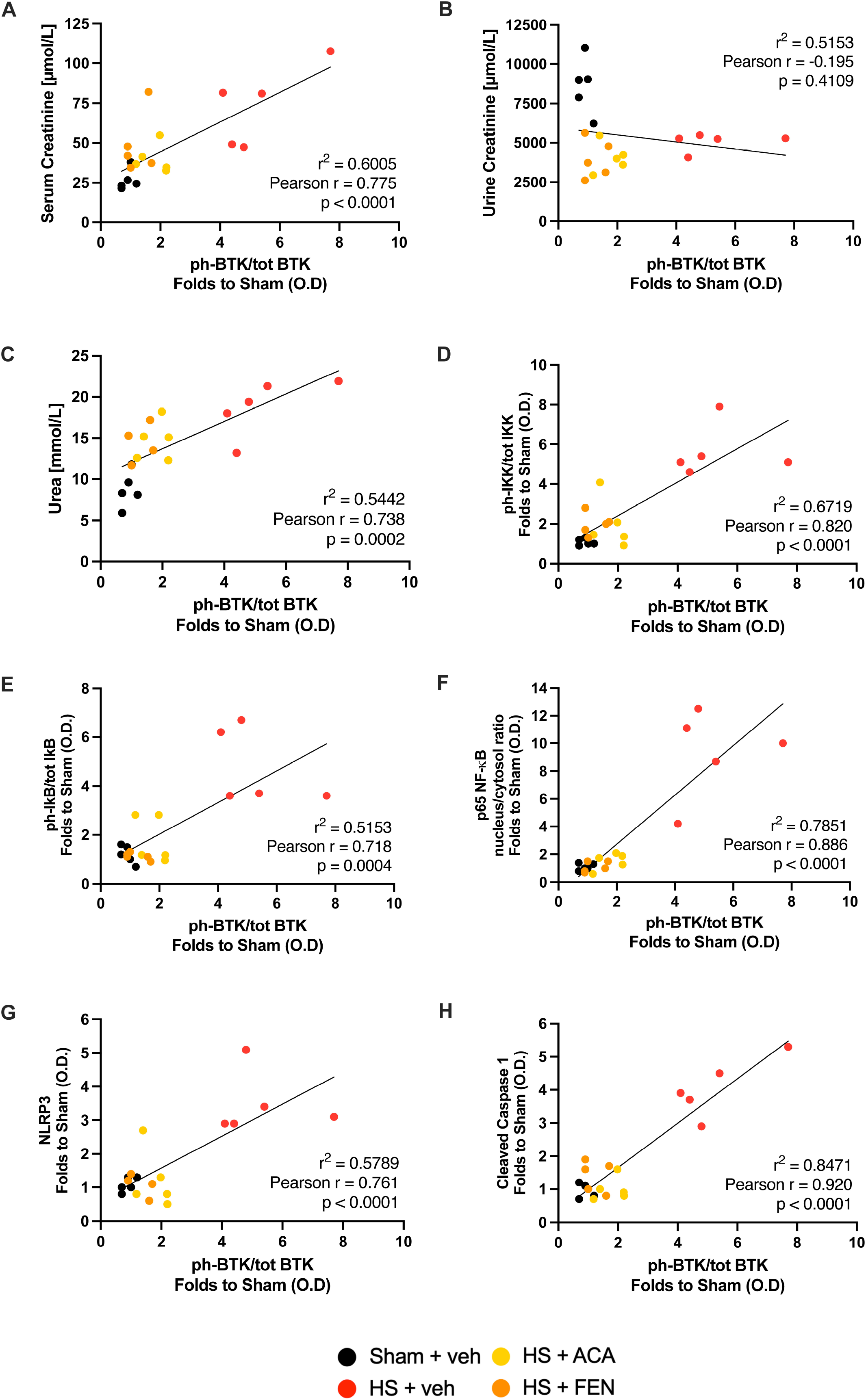
BTK activation correlates with renal dysfunction, NF-κB and NLRP3 activation in an acute HS model. Linear regression analysis of (**A**) phosphorylation of BTK at Tyr^223^ vs. serum creatinine, (**B**) phosphorylation of BTK at Tyr^223^ vs. urine creatinine, (**C**) phosphorylation of BTK at Tyr^223^ vs. serum urea, (**D**) phosphorylation of BTK at Tyr^223^ vs. phosphorylation of IKKα/β at Ser^176/180^, (**E**) phosphorylation of BTK at Tyr^223^ vs. phosphorylation of IκBα at Ser^32/36^ (**F**) phosphorylation of BTK at Tyr^223^ vs. translocation of p65 (**G**) phosphorylation of BTK at Tyr^223^ vs. expression of NLRP3 and (**H**) phosphorylation of BTK at Tyr^223^ vs. expression of the cleaved form of caspase 1. Data are expressed as raw individual values of five animals per group. Statistical analysis was performed using simple linear regression to calculate the r^2^ value, Pearson correlation coefficient test to calculate the r value and a two-tailed t-test to calculate the p-value. *p<0.05 denoted statistical significance.

### Treatment with acalabrutinib improves HS-induced circulatory failure in a chronic HS model

When comparing acalabrutinib and fenebrutinib, both inhibitors were equally efficacious in the acute HS model but acalabrutinib has the advantage of being FDA approved; hence was investigated in a chronic HS model. Having demonstrated treatment with acalabrutinib improved blood pressure in an acute HS rat model, we wished to determine whether acalabrutinib would still be effective in a HS model in which the resuscitation period is prolonged to 24 h. When compared to sham-operated rats, HS-rats treated with vehicle had significantly lower values of MAP recorded 24 h after the onset of resuscitation (p<0.05; Figure 6A); highlighting that either cardiac dysfunction or excessive hypotension^23^ was still present. In contrast, the MAP values of HS-rats treated with acalabrutinib upon resuscitation were significantly higher even at 24 h after the onset of resuscitation [when compared with those of vehicle treated rats (p<0.05; Figure 6A)]. Administration of acalabrutinib to sham-operated rats had no significant effect on MAP (p>0.05; Figure 6A). There were no significant differences in HR between any of the four groups investigated (p>0.05; Figure 6B).

**Figure 6:**
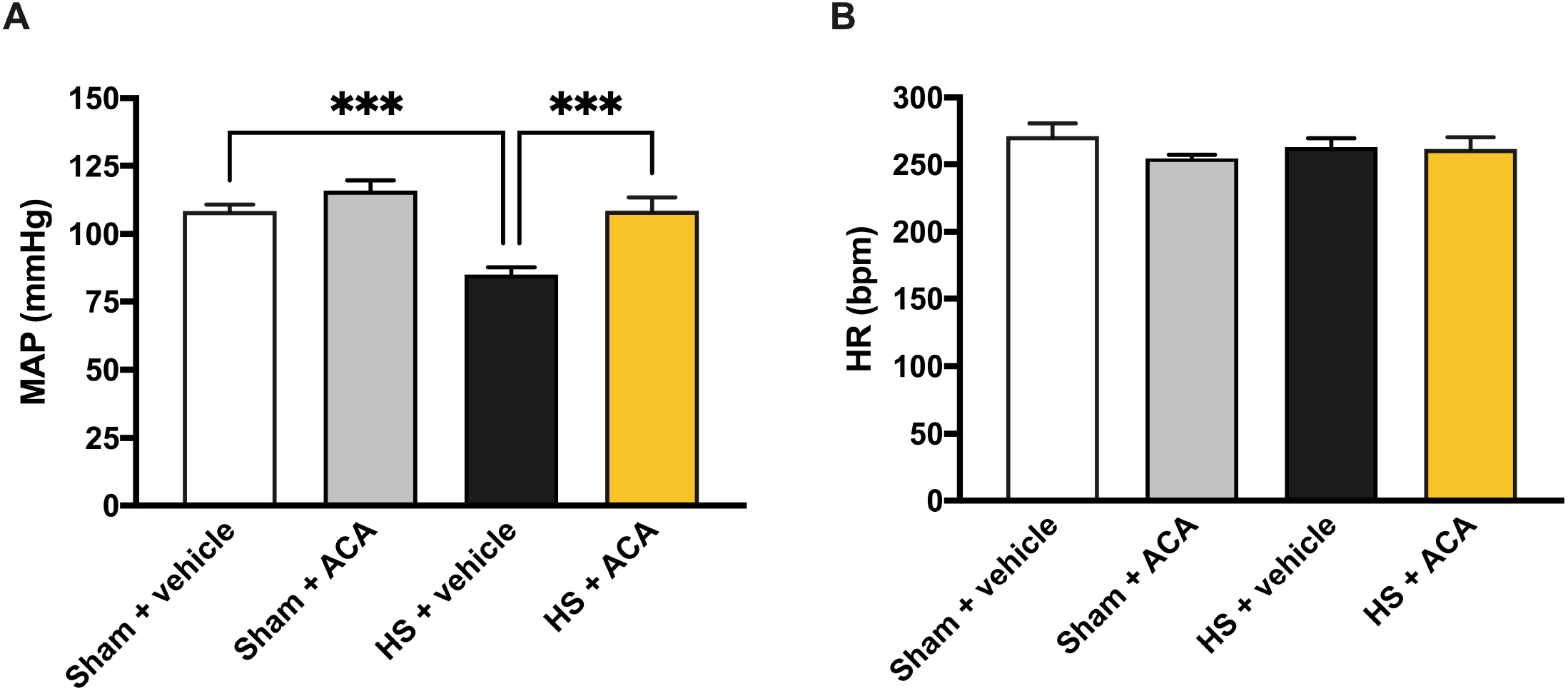
Treatment with acalabrutinib improves HS-induced circulatory failure in a chronic HS model. (**A**) Mean arterial pressure (MAP) and (**B**) heart rate (HR) were measured 24 h post resuscitation for vehicle and acalabrutinib (ACA) treated rats. Sham-operated rats were used as control. Data are expressed as mean ± SEM of 9-10 animals per group. Statistical analysis was performed using one-way ANOVA followed by a Bonferroni’s *post-hoc* test. *p<0.05 denoted statistical significance.

### Treatment with acalabrutinib attenuates HS-induced organ damage in a chronic HS model

Having shown that treatment with acalabrutinib ameliorated the MODS associated with HS in an acute HS rat model, we examined whether this effect was sustained when the resuscitation period was extended to 24 h. As with the acute HS model, when compared to sham-operated rats, rats subjected to chronic HS and treated with vehicle displayed significant increases in serum urea (p<0.05; Figure 7A) and creatinine (p<0.05; Figure 7B); indicating the development of renal dysfunction. When compared to sham-operated rats, vehicle treated HS-rats exhibited significant increases in ALT (p<0.05; Figure 7C) and AST (p<0.05; Figure 7D) indicating the development of hepatic injury, whilst the significant increase in LDH (p<0.05; Figure 7E) confirmed general tissue injury. Treatment of HS-rats with acalabrutinib significantly attenuated the hepatic and general tissue injury caused by HS as shown by the decrease in serum parameter values (p<0.05; Figures 7C-E). Treatment with acalabrutinib had no significant effect on renal dysfunction (p>0.05; Figures 7A-B). Administration of acalabrutinib to sham-operated rats had no significant effect on any of the parameters measured (p>0.05; Figures 7A-E).

**Figure 7:**
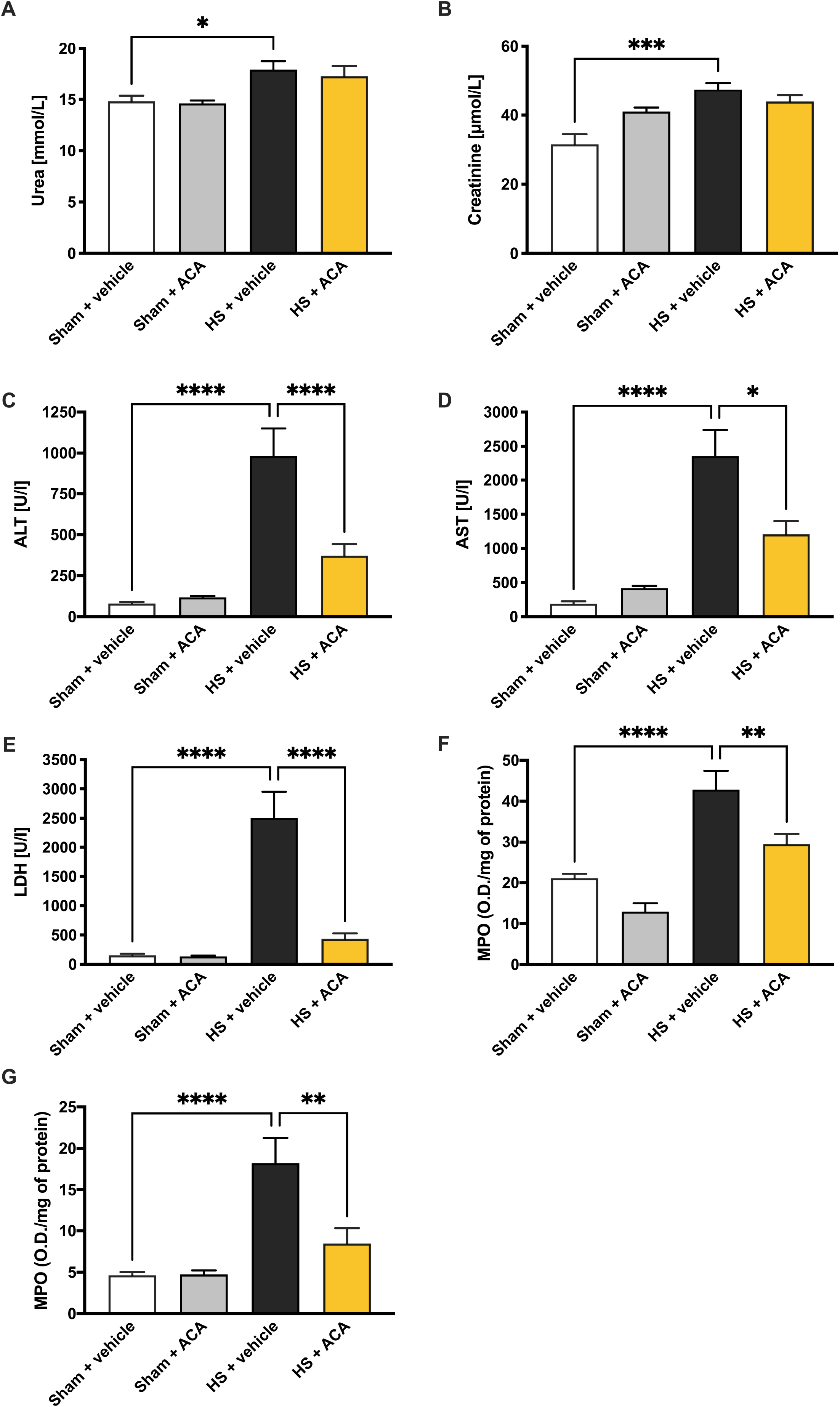
Treatment with acalabrutinib attenuates HS-induced organ damage and myeloperoxidase activity in a chronic HS model. Rats were subjected to hemorrhagic shock (HS) and 24 h after resuscitation, levels of serum (**A**) urea, (**B**) creatinine, (**C**) alanine aminotransferase (ALT), (**D**) aspartate aminotransferase (AST) and (**E**) lactate dehydrogenase (LDH) and myeloperoxidase (MPO) activity in the (**F**) lung and (**G**) liver were determined for vehicle and acalabrutinib (ACA) treated rats. Sham-operated rats were used as control. Data are expressed as mean ± SEM of 9-10 animals per group. Statistical analysis was performed using one-way ANOVA followed by a Bonferroni’s *post-hoc* test. *p<0.05 denoted statistical significance.

### Treatment with acalabrutinib reduces myeloperoxidase activity in a chronic HS model

We next determined myeloperoxidase (MPO) activity in the lung and liver of rats subjected to HS as an indicator of neutrophil infiltration. When compared to sham-operated rats, HS-rats treated with vehicle showed a significant increase in MPO activity in the lung (p<0.05; Figure 7F) and liver (p<0.05; Figure 7G). Treatment with acalabrutinib in HS-rats significantly attenuated the rises in pulmonary (Figure 7F) and hepatic (Figure 7G) MPO activity (p<0.05), suggesting reduced neutrophil recruitment and inflammation. Administration of acalabrutinib to sham-operated rats had no significant effect on both pulmonary and hepatic MPO activity (p>0.05; Figures 7F-G).

## DISCUSSION

This study reports for the first time that inhibition of BTK activity attenuates organ injury/dysfunction and circulatory failure in acute (Figures 2 and 3) and chronic (Figures 6 and 7) rat models of HS. Having shown that BTK gene expression is significantly elevated in leukocytes of trauma patients (Figure 1), we used a reverse translational approach to investigate whether pharmacological intervention with acalabrutinib and fenebrutinib, ameliorates the MODS associated with HS in a well-established rat model. Administration of either an irreversible (acalabrutinib) or a reversible (fenebrutinib) inhibitor of BTK activity significantly attenuated the fall in blood pressure caused by HS (Figures 2 acute and 6 chronic). Thus, BTKi reduces the delayed vascular decompensation which is (in part) secondary to the expression of inducible nitric oxide synthase^23^. Moreover, BTKi significantly attenuated the renal dysfunction, hepatic injury and neuromuscular injury caused by HS (Figures 3 acute and 7 chronic). This is consistent with published studies which demonstrated a reduction in disease severity following BTK inhibition in animal models of lupus nephritis^24,25^, warm hepatic ischemia and reperfusion^26^, acute lung injury^27–29^ and sepsis^9^.

What, then, are the mechanisms by which BTKi attenuate HS-associated organ injury/dysfunction? Western blot analysis revealed that acute HS resulted in a significant increase in BTK activity in the injured kidney (Figure 4). Most notably, this activation of BTK positively correlated with serum creatinine and urea (Figure 5), indicating that activation of BTK is associated with the renal dysfunction in hemorrhagic shock. Indeed, inhibition of BTK activity with acalabrutinib or fenebrutinib in the kidney of shocked animals decreases the renal dysfunction in hemorrhagic shock, suggesting that activation of BTK plays a pivotal role in the pathophysiology of the renal dysfunction in hemorrhagic shock. Other research groups have shown a positive correlation between BTK expression and creatinine in patients with IgA nephropathy^30^ and diabetic nephropathy^31^. Our finding is the first in the field of trauma and hemorrhagic shock.

There is good evidence that BTK activation precedes the activation of NF-κB through TLR signaling^32^. Indeed, trauma results in elevated NF-κB translocation to the nucleus^19–22^. Inhibition of BTK activity with acalabrutinib or fenebrutinib reduced the activation of NF-κB in the kidney (Figure 4). We also found a significant positive correlation between the activation of BTK and the phosphorylation of IKKα/β at Ser^176/180^, phosphorylation of IκBα at Ser^32/36^ and translocation of p65 (Figure 5). This may suggest that inhibiting the activation of NF-κB contributes to the observed beneficial effects of acalabrutinib and fenebrutinib in trauma-hemorrhage; as NF-κB activation is involved in the regulation of immunological and inflammatory responses. Activation of NF-κB drives the formation of several pro- and anti-inflammatory mediators which include cytokines, chemokines and enzymes^33^. As part of a positive feedback loop, these mediators can activate NF-κB and its upstream signaling components, further amplifying and perpetuating the inflammatory responses mediated by NF-κB which can lead to increased endothelial permeability, tissue hypoperfusion/hypoxia, tissue injury and ultimately MODS^11^. It should be noted that the inhibition of BTK activity in CLP-sepsis also reduced activation of NF-κB^9^.

There is also good evidence that BTK activation influences the assembly and activation of the NLRP3 inflammasome in rodents and humans^34–36^. NLRP3 inflammasome activation drives the production of IL-1β which plays a crucial role in the systemic inflammation and/or organ dysfunction in trauma^22^. Inhibition of BTK activity with acalabrutinib or fenebrutinib reduces both the assembly and subsequent activation of the NLRP3 inflammasome in the kidney (Figure 4). We also discovered a significant positive correlation between the activation of BTK and both the activation of NLRP3 and caspase 1 (Figure 5). This may suggest that inhibiting the activation of the NLRP3 inflammasome contributes to the observed protective effects of acalabrutinib and fenebrutinib in trauma-hemorrhage by lowering the pro-inflammatory effects of IL-1β and resulting tissue inflammation. This is due to IL-1β not only directly causing inflammation but also stimulating the expression of several pro-inflammatory adhesion molecules and cytokines which further exacerbates the inflammatory response^37^. It should be noted that the inhibition of BTK activity in CLP-sepsis also reduced activation of NLRP3 and cleavage of pro-caspase 1 to caspase 1^9^.

The sterile inflammation caused by HS drives leukocyte recruitment to the tissues and is secondary to the activation of NF-κB and NLRP3 and their transcriptional regulation of pro-inflammatory cytokines^38–40^. Moreover, the expression of adhesion molecules present on leukocytes and endothelial cells is regulated by NF-κB and permits leukocyte extravasation from the circulation to the site of injury^41^. As neutrophils play a key role in HS-associated pulmonary and hepatic inflammation, we evaluated the degree of neutrophil recruitment (measured as MPO activity) in the lung and liver^42,43^. HS resulted in a significant increase in both pulmonary and hepatic MPO activity which was attenuated by the treatment of HS-rats with acalabrutinib (Figure 7). Following ICAM-1 binding on endothelial cells and subsequent tissue recruitment, neutrophils undergo degranulation and release pro-inflammatory mediators^44,45^. These mediators (e.g. cytokines, reactive oxygen species and MPO) can contribute to inflammation by exerting direct cytotoxic cellular effects at the local site; subsequently leading to organ dysfunction and potential mortality following HS and resuscitation^3,46^. As we have demonstrated in the acute HS model that treatment with BTKi attenuates both NF-κB and NLRP3 activation, it can be implied that the decreased recruitment of neutrophils and lowered inflammation is secondary to this reduced activation of NF-κB and NLRP3.

Our results and conclusions are supported by findings in patient cohorts with either COVID-19 or sepsis where BTK has been proposed to play a role in disease pathogenesis; both diseases featuring the hallmarks of excessive systemic inflammation similar to those seen in trauma-associated MODS^15,47^. The beneficial effects of acalabrutinib in COVID-19 patients, as measured by the determination of biomarkers of inflammation, oxygenation and clinical status, imply that BTK activation plays a role in the pathology^15^. Whilst Parnell and colleagues did not investigate pharmacological intervention with BTKi in patients with sepsis, reanalysis of microarray data (Gene Expression Omnibus Dataset Number GDS4971) revealed an increased expression (in whole blood) of BTK in septic non-survivors compared to septic survivors; highlighting the potential for BTK to be a predictor of mortality^10^.

## LIMITATIONS OF THE STUDY

Although acalabrutinib and fenebrutinib displayed some striking, beneficial effects in the HS models, there are study limitations which should be considered. Further long-term survival experiments are needed to verify that the observed early reduction in MODS does, indeed, translate to improved outcome and ultimately reduced mortality. In our study, organ injury and dysfunction were used as surrogate markers for mortality (as the determination of mortality is not allowed by our respective ethics and/or Home Office licenses). Therefore, care must be taken when interpreting our pre-clinical results and extrapolating them to the clinical scenario. In addition, future studies in larger animals and/or higher species may be useful to confirm efficacy and to further investigate the mechanism of action (e.g. blood gas analysis and microcirculatory effects) of BTKi in HS. As BTK is primarily expressed in cells of hematopoietic lineage (except T-lymphocytes, plasma cells and natural killer cells), it is likely that the increase in BTK activity measured in the kidneys of rats with HS was secondary to the infiltration of these organs by invading immune cells rather than an increase in parenchymal renal tissue (where no expression of BTK has been reported). Therefore, it is possible that inhibition of BTK leads to decreased recruitment of leukocytes into the kidney (as a result of reduced NF-κB and NLRP3 activation) and subsequently to lower levels of BTK activation in the kidney. Furthermore, clinical studies with large cohorts of trauma patients are needed to robustly examine the relationship between BTK activity and clinical outcomes in humans.

## CONCLUSIONS

In conclusion, we report here for the first time that treatment with either the irreversible BTKi acalabrutinib or the reversible BTKi fenebrutinib reduces the organ injury/dysfunction and circulatory failure caused by severe hemorrhage in the rat; highlighting a role of BTK in disease pathogenesis. Moreover, experimental trauma-hemorrhage results in a significant upregulation of BTK in the kidney. Administration of either BTKi subsequently attenuates the degree of activation of BTK as well as the activation of NF-κB and the NLRP3 inflammasome (measured in the kidney), both of which are key drivers of local and systemic inflammation. Notably, no significant differences were found between the two structurally and mechanistically different inhibitors, suggesting that the observed beneficial effects in experimental trauma-hemorrhage are most likely due to a drug class related effect. Thus, we propose that BTKi may be repurposed for the use in trauma patients to lower the organ injury and inflammation caused by severe hemorrhage and resuscitation.

## ABBREVIATIONS

ALT: alanine aminotransferase
AST: aspartate aminotransferase
BTK: Bruton’s tyrosine kinase
BTKi: Bruton’s tyrosine kinase inhibitors
CK: creatine kinase
DAMP: damage-associated molecular pattern
FDA: Food and Drug Administration
HR: heart rate
HS: hemorrhagic shock
I/R: ischemia-reperfusion
LDH: lactate dehydrogenase
MAP: mean arterial pressure
MODS: multiple organ dysfunction syndrome
MPO: myeloperoxidase

## LIST OF SUPPLEMENTAL DIGITAL CONTENT

Supplemental digital content: BTK in Trauma Supplemental.pdf

## REFERENCES

1. World Health Organization. Injuries and Violence: The Facts 2014.; 2014. Accessed May 6, 2021. www.who.int/healthinfo/global_burden_disease/projections/en/

2. Curry N, Hopewell S, Dorée C, Hyde C, Brohi K, Stanworth S. The acute management of trauma hemorrhage: A systematic review of randomized controlled trials. Critical Care. 2011;15(2):1–10. doi:10.1186/cc10096

3. Dewar D, Moore FA, Moore EE, Balogh Z. Postinjury multiple organ failure. Injury. 2009;40(9):912–918. doi:10.1016/j.injury.2009.05.024

4. Lord JM, Midwinter MJ, Chen YF, et al. The systemic immune response to trauma: An overview of pathophysiology and treatment. The Lancet. 2014;384(9952):1455–1465. doi:10.1016/S0140-6736(14)60687-5

5. Vetrie D, Vořechovský I, Sideras P, et al. The gene involved in X-linked agammaglobulinaemia is a member of the src family of protein-tyrosine kinases. Nature. 1993;361(6409):226–233. doi:10.1038/361226a0

6. Mohamed AJ, Yu L, Bäckesjö CM, et al. Bruton’s tyrosine kinase (Btk): Function, regulation, and transformation with special emphasis on the PH domain. Immunological Reviews. 2009;228(1):58–73. doi:10.1111/j.1600-065X.2008.00741.x

7. Weber ANR, Bittner Z, Liu X, Dang TM, Radsak MP, Brunner C. Bruton’s tyrosine kinase: An emerging key player in innate immunity. Frontiers in Immunology. 2017;8(NOV):1454. doi:10.3389/fimmu.2017.01454

8. Xiao W, Mindrinos MN, Seok J, et al. A genomic storm in critically injured humans. The Journal of Experimental Medicine. 2011;208(13):2581. doi:10.1084/JEM.20111354

9. O’Riordan CE, Purvis GSD, Collotta D, et al. Bruton’s Tyrosine Kinase Inhibition Attenuates the Cardiac Dysfunction Caused by Cecal Ligation and Puncture in Mice. Frontiers in Immunology. 2019;10:2129. doi:10.3389/fimmu.2019.02129

10. O’Riordan CE, Purvis GSD, Collotta D, et al. X-Linked Immunodeficient Mice With No Functional Bruton’s Tyrosine Kinase Are Protected From Sepsis-Induced Multiple Organ Failure. Frontiers in Immunology. 2020;11:581758. doi:10.3389/fimmu.2020.581758

11. Liu SF, Malik AB. NF-κB activation as a pathological mechanism of septic shock and inflammation. American Journal of Physiology - Lung Cellular and Molecular Physiology. 2006;290(4). doi:10.1152/ajplung.00477.2005

12. Coldewey SM, Rogazzo M, Collino M, Patel NSA, Thiemermann C. Inhibition of IκB kinase reduces the multiple organ dysfunction caused by sepsis in the mouse. Disease Models & Mechanisms. 2013;6(4):1031. doi:10.1242/DMM.012435

13. Sordi R, Chiazza F, Johnson FL, et al. Inhibition of IκB kinase attenuates the organ injury and dysfunction associated with hemorrhagic shock. Molecular Medicine. 2015;21(1):563–575. doi:10.2119/molmed.2015.00049

14. Nicolson PLR, Welsh JD, Chauhan A, Thomas MR, Kahn ML, Watson SP. A rationale for blocking thromboinflammation in COVID-19 with Btk inhibitors. Platelets. 2020;31(5):685–690. doi:10.1080/09537104.2020.1775189

15. Roschewski M, Lionakis MS, Sharman JP, et al. Inhibition of Bruton tyrosine kinase in patients with severe COVID-19. Science Immunology. 2020;5(48). doi:10.1126/SCIIMMUNOL.ABD0110

16. Soresina A, Moratto D, Chiarini Marco, et al. Two X-linked agammaglobulinemia patients develop pneumonia as COVID-19 manifestation but recover. Pediatr Allergy Immunol. 2020;00:1–5. doi:10.1111/pai.13263

17. Burger JA. Bruton Tyrosine Kinase Inhibitors: Present and Future. Cancer Journal (United States). 2019;25(6):386–393. doi:10.1097/PPO.0000000000000412

18. Liu X, Zhang J, Han W, et al. Inhibition of BTK protects lungs from trauma-hemorrhagic shock-induced injury in rats. Molecular Medicine Reports. 2017;16(1):192–200. doi:10.3892/mmr.2017.6553

19. Sordi R, Chiazza F, Collotta D, et al. Resolvin D1 Attenuates the Organ Injury Associated With Experimental Hemorrhagic Shock. Annals of Surgery. 2021;273(5):1012–1021. doi:10.1097/sla.0000000000003407

20. Sordi R, Nandra KK, Chiazza F, et al. Artesunate protects against the organ injury and dysfunction induced by severe hemorrhage and resuscitation. Annals of Surgery. Published online 2017. doi:10.1097/SLA.0000000000001664

21. Yamada N, Martin LB, Zechendorf E, et al. Novel Synthetic, Host-defense Peptide Protects Against Organ Injury/Dysfunction in a Rat Model of Severe Hemorrhagic Shock. Annals of Surgery. Published online 2018. doi:10.1097/SLA.0000000000002186

22. Martin L, Patel NM, 2# M, et al. The inhibition of Macrophage Migration Inhibitory Factor by ISO-1 attenuates trauma-induced multi organ dysfunction in rats. medRxiv. Published online April 29, 2021:2021.04.28.21255719. doi:10.1101/2021.04.28.21255719

23. Thiemermann C, Szabo C, Mitchell JA, Vane JR. Vascular hyporeactivity to vasoconstrictor agents and hemodynamic decompensation in hemorrhagic shock is mediated by nitric oxide. Proceedings of the National Academy of Sciences of the United States of America. 1993;90(1):267–271. doi:10.1073/pnas.90.1.267

24. Chalmers SA, Doerner J, Bosanac T, et al. Therapeutic Blockade of Immune Complex-Mediated Glomerulonephritis by Highly Selective Inhibition of Bruton’s Tyrosine Kinase. Scientific Reports. 2016;6. doi:10.1038/srep26164

25. Chalmers SA, Glynn E, Garcia SJ, et al. BTK inhibition ameliorates kidney disease in spontaneous lupus nephritis. Clinical Immunology. 2018;197:205–218. doi:10.1016/j.clim.2018.10.008

26. Palumbo T, Nakamura K, Lassman C, et al. Bruton Tyrosine Kinase Inhibition Attenuates Liver Damage in a Mouse Warm Ischemia and Reperfusion Model. Transplantation. 2017;101(2):322–331. doi:10.1097/TP.0000000000001552

27. Krupa A, Fol M, Rahman M, et al. Silencing bruton’s tyrosine kinase in alveolar neutrophils protects mice from LPS/immune complex-induced acute lung injury. American Journal of Physiology - Lung Cellular and Molecular Physiology. 2014;307(6):L435–L448. doi:10.1152/ajplung.00234.2013

28. Zhou P, Ma B, Xu S, et al. Knockdown of Burton’s tyrosine kinase confers potent protection against sepsis-induced acute lung injury. Cell biochemistry and biophysics. 2014;70(2):1265–1275. doi:10.1007/s12013-014-0050-1

29. Florence JM, Krupa A, Booshehri LM, Davis SA, Matthay MA, Kurdowska AK. Inhibiting bruton’s tyrosine kinase rescues mice from lethal influenza-induced acute lung injury. American Journal of Physiology - Lung Cellular and Molecular Physiology. 2018;315(1):L52–L58. doi:10.1152/ajplung.00047.2018

30. Wei J, Wang Y, Qi X, Wu Y. Enhanced Bruton’s tyrosine kinase activity in the kidney of patients with IgA nephropathy. International Urology and Nephrology. 2021;53(7):1399. doi:10.1007/S11255-020-02733-2

31. Zhao J, Chen J, Li Y-Y, Xia L-L, Wu Y-G. Bruton’s tyrosine kinase regulates macrophage-induced inflammation in the diabetic kidney via NLRP3 inflammasome activation. International Journal of Molecular Medicine. 2021;48(3). doi:10.3892/IJMM.2021.5010

32. Jefferies CA, Doyle S, Brunner C, et al. Bruton’s tyrosine kinase is a Toll/interleukin-1 receptor domain-binding protein that participates in nuclear factor κB activation by toll-like receptor 4. Journal of Biological Chemistry. 2003;278(28):26258–26264. doi:10.1074/jbc.M301484200

33. Senftleben U, Karin M. The IKK/NF-kappa B pathway. Crit Care Med. 2002;30:S18–S26. Accessed May 11, 2021. https://pubmed.ncbi.nlm.nih.gov/11782557/

34. Ito M, Shichita T, Okada M, et al. Bruton’s tyrosine kinase is essential for NLRP3 inflammasome activation and contributes to ischaemic brain injury. Nature Communications. 2015;6(1):1–11. doi:10.1038/ncomms8360

35. Liu X, Pichulik T, Wolz OO, et al. Human NACHT, LRR, and PYD domain–containing protein 3 (NLRP3) inflammasome activity is regulated by and potentially targetable through Bruton tyrosine kinase. Journal of Allergy and Clinical Immunology. 2017;140(4):1054-1067.e10. doi:10.1016/j.jaci.2017.01.017

36. Bittner ZA, Liu X, Shankar S, et al. BTK operates a phospho-tyrosine switch to regulate NLRP3 inflammasome activity. bioRxiv. Published online June 25, 2020:864702. doi:10.1101/864702

37. Dinarello CA. Biologic basis for interleukin-1 in disease. Blood. 1996;87(6):2095–2147. doi:10.1182/blood.v87.6.2095.bloodjournal8762095

38. Chen CC, Manning AM. Transcriptional regulation of endothelial cell adhesion molecules: A dominant role for NF-κB. Agents and Actions Supplements. 1995;47:135–141. doi:10.1007/978-3-0348-7343-7_12

39. Collins T, Read MA, Neish AS, Whitley MZ, Thanos D, Maniatis T. Transcriptional regulation of endothelial cell adhesion molecules: NF-κB and cytokine-inducible enhancers. The FASEB Journal. 1995;9(10):899–909.

40. Campbell SJ, Anthony DC, Oakley F, et al. Hepatic Nuclear Factor κB Regulates Neutrophil Recruitment to the Injured Brain. Journal of Neuropathology & Experimental Neurology. 2008;67(3):223–230. doi:10.1097/NEN.0B013E3181654957

41. Hayden MS, Ghosh S. NF-κB in immunobiology. Cell Research 2011 21:2. 2011;21(2):223–244. doi:10.1038/cr.2011.13

42. Botha AJ, Moore FA, Moore EE, Kim FJ, Banerjee A, Peterson VM. Postinjury neutrophil priming and activation: An early vulnerable window. Surgery. 1995;118(2):358–365. doi:10.1016/S0039-6060(05)80345-9

43. Partrick D, Moore F, Moore E, Barnett Jr C, Silliman C. Neutrophil priming and activation in the pathogenesis of postinjury multiple organ failure. New Horiz. 1996;4(2):194–210. Accessed May 12, 2021. https://pubmed.ncbi.nlm.nih.gov/8774796/

44. Sauaia A, Moore FA, Moore EE. Postinjury Inflammation and Organ Dysfunction. Critical Care Clinics. 2017;33(1):167–191. doi:10.1016/j.ccc.2016.08.006

45. Olanders K, Sun Z, Börjesson A, et al. The effect of intestinal ischemia and reperfusion injury on ICAM-1 expression, endothelial barrier function, neutrophil tissue influx, and protease inhibitor levels in rats. Shock. 2002;18(1):86–92. doi:10.1097/00024382-200207000-00016

46. Zhang Y, Zhang J, Korff S, Ayoob F, Vodovotz Y, Billiar TR. Delayed neutralization of interleukin 6 reduces organ injury, selectively suppresses inflammatory mediator, and partially normalizes immune dysfunction following trauma and hemorrhagic shock. Shock (Augusta, Ga). 2014;42(3):218–227. doi:10.1097/SHK.0000000000000211

47. Parnell GP, Tang BM, Nalos M, et al. Identifying key regulatory genes in the whole blood of septic patients to monitor underlying immune dysfunctions. Shock. 2013;40(3):166–174. doi:10.1097/SHK.0b013e31829ee604

