## Supplemental Methods/Figures for "Inhibition of Bruton’s tyrosine kinase activity attenuates trauma-induced multiple organ dysfunction in rats"

### *Acute Hemorrhagic Shock Model*

Rats were anesthetized with sodium thiopentone (120 mg/kg i.p. initially and 10 mg/kg i.v. for maintenance as needed). Cannulation with polyethylene catheters (Smiths Medical International Ltd., Kent, UK) of the trachea for facilitation of spontaneous breathing (internal diameter [ID] 1.67 mm), left femoral artery for recording of the mean arterial pressure (MAP) (ID 0.40 mm), left carotid artery for blood withdrawal (ID 0.58 mm) and right jugular vein for fluid and drug administration (ID 0.40 mm) was performed. To prevent tissue desiccation, swabs moistened with saline were placed over the surgery incision sites. Body temperature was monitored by a rectal probe thermometer and maintained at 37°C ± 0.3°C by means of a homoeothermic blanket system (Harvard Apparatus). Upon the completion of surgery, the MAP was allowed to stabilize for 15 min. Blood was then withdrawn (up to 1 mL/min into heparinized syringes containing 100 IU/mL heparin mixed with normal saline) through the cannula inserted in the carotid artery in order to achieve a fall in MAP to 35 ± 5 mmHg, which was recorded with a pressure transducer (attached to the femoral artery cannula, 844-31 Memscap, Durham, USA) and coupled to a PowerLab 8/30 data acquisition system (AD Instruments Pty Ltd., Castle Hill, Australia). Thereafter, MAP was maintained at 35 ± 5 mmHg for a period of 90 min either by further withdrawal of blood during the compensation phase or administration of the shed blood during the decompensation phase. At 90 min after initiation of hemorrhage (or when 25% of the shed blood had to be re-injected to sustain MAP at 35 ± 5 mmHg), resuscitation through the jugular vein was performed with the remaining shed blood (mixed with 100 IU/mL heparinized saline) over a period of 5 min plus a volume of Ringer's lactate identical to the volume of shed blood. Treatment or vehicle was also administered intravenously. One hour after resuscitation, an infusion of Ringer's lactate (1.5

mL/kg/h) was started as fluid replacement and it was maintained throughout the experiment for a total of 3 h. Under deep anesthesia, the heart was removed to terminate the experiment 4 h after resuscitation. Sham-operated rats were used as control and underwent identical surgical procedures, but without hemorrhage or resuscitation.

Rats were treated with either acalabrutinib (3 mg/kg), fenebrutinib (3 mg/kg) or its vehicle (5% DMSO + 95% Ringer's lactate) intravenously as a bolus treatment immediately after resuscitation. This dose of acalabrutinib was based on the dose used in studies conducted by the Thiemermann group[9]. The dose of fenebrutinib was chosen to match that of acalabrutinib.

*Sample Collection - Acute Hemorrhagic Shock Model*

Rats remained anesthetized with sodium thiopentone (120 mg/kg i.p.) before sacrifice. Up to 5 mL of blood was taken from the carotid artery via the inserted cannula into a non-heparinized 5 mL syringe and immediately decanted into 1.1 mL serum gel tubes (Sarstedt, Germany). The blood was centrifuged (10,000 g for 5 min) to obtain the serum, which was subsequently stored at -80 °C until analysis. Organs (heart, lungs, liver, spleen and kidneys) were excised of which one section was snap frozen in liquid nitrogen and stored at -80 °C, and another section was placed in 10 % formalin for 24-48 h; followed by transfer to 70 % ethanol until further analysis. All organ injury/dysfunction parameters (urea, creatinine, alanine aminotransferase [ALT], aspartate aminotransferase [AST], creatine kinase [CK], amylase and lactate dehydrogenase [LDH]) in the serum were measured in a blinded fashion by a clinical pathology diagnostic laboratory (MRC Harwell Institute, Oxfordshire, UK).

*Chronic Hemorrhagic Shock Model*

At 15 min prior to anesthesia, analgesia with tramadol (10 mg/kg i.p.) was administered. Rats were then anesthetized with ketamine-xylazine (ketamine, 100 mg/kg; xylazine, 10 mg/kg i.m. initially and 100 µL ketamine i.p. for maintenance as needed). Cannulation with polyethylene catheters of the left femoral artery and left femoral vein was performed. Body temperature was monitored by a digital ear thermometer and maintained at 36.5 °C ± 0.5 °C by means of a homoeothermic blanket system (Harvard Apparatus). Upon completion of surgery, the MAP was allowed to stabilize for 15 min. Blood was then withdrawn (up to 1 mL/min into heparinized syringes containing 100 IU/mL heparin mixed with normal saline) through the cannula inserted in the femoral artery in order to achieve a fall in MAP to 40 ± 2 mmHg, which was recorded with a pressure transducer (attached to the femoral artery cannula) and coupled to a PowerLab 8/30 (AD Instruments Pty Ltd., Castle Hill, Australia). Thereafter, MAP was maintained at 40 ± 2 mmHg for a period of 90 min either by further withdrawal of blood or administration of the shed blood. At 90 min after initiation of hemorrhage (or when 25 % of the shed blood had to be re-injected to sustain MAP at 40 ± 2 mmHg), resuscitation through the femoral vein was performed with the remaining shed blood over a period of 5 min plus 1.5 mL/kg Ringer's lactate. An initial bolus of acalabrutinib treatment (1.5 mg/kg) or vehicle was then administered intraperitoneally. Sham-operated rats were used as control and underwent identical surgical procedures, but without hemorrhage or resuscitation. At 20 min after resuscitation was completed, the catheters were removed, the femoral vessels were ligated, and the incision was closed with sutures. Rats were allowed to recover from the anesthesia and given tramadol (5 mg/kg i.p.) 12 h later in addition to a second dose of acalabrutinib (1.5 mg/kg) or vehicle (i.p.).

*Western Blot Analysis*

Semi-quantitative immunoblot analysis were carried out in kidney tissue samples as previously described[21]. Briefly, kidney samples were homogenized in buffer and centrifuged (1320 g, 5 mins, 4 °C). To obtain the cytosolic fraction, supernatants were centrifuged (16,125 g, 4 °C, 40 mins). The pelleted nucleoli were resuspended in extraction buffer and centrifuged (16,125 g, 20 mins, 4 °C). Protein content was determined on both nuclear and cytosolic extracts using bicinchoninic acid (BCA) protein assay (Thermo Fisher Scientific Inc, Rockford, IL). Proteins were separated by 8% sodium dodecyl sulfate polyacrylamide (SDS-PAGE) gel electrophoresis and electrotransferred to polyvinylidene difluoride (PVDF) membrane. After blocking (1 hr in 10% dry milk solution), membranes were incubated with primary antibodies in 5% blocking solution overnight [1:1000 rabbit anti-total BTK, 1:1000 rabbit anti-NF-κB, 1:1000 rabbit anti-IKKβ, 1:1000 rabbit anti-Ser$^{176/180}$ IKKα/β, 1:1000 mouse anti-Ser$^{32/36}$ IκBα, 1:1000 mouse anti-total IκBα (from Cell Signaling), 1:1000 rabbit anti-Tyr$^{223}$-BTK, 1:1000 rabbit anti-NLRP3 inflammasome (from Abcam), 1:1000 mouse anti-caspase 1 (p20) (from Adipogen)] followed by incubation with appropriate HRP-conjugated secondary antibodies. Proteins were detected with an ECL detection system and quantified by densitometry using analytic software (Quantity-One; Bio-Rad, Hercules, CA). Results were normalized with respect to densitometric values of tubulin for cytosolic proteins or histone H3 for nuclear proteins.

**SUPPLEMENTAL FIGURES**

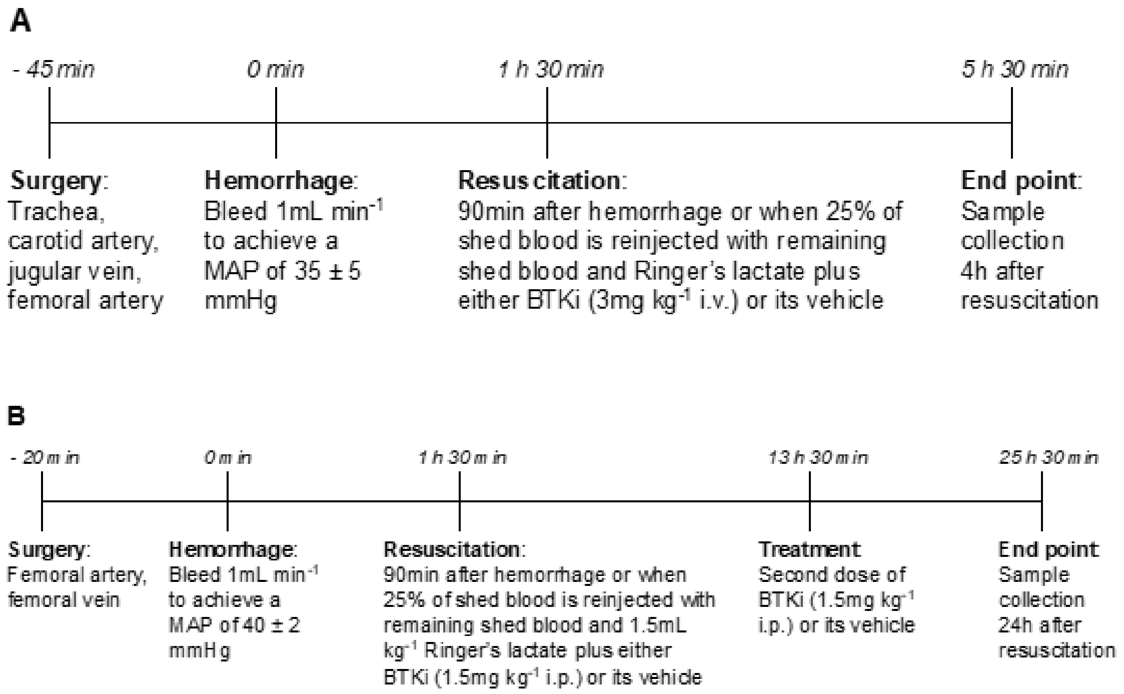


**Supplemental Figure 1: Schematic representations of the acute and chronic HS models.** The experimental procedures at each stage of the (**A**) acute and (**B**) chronic HS models are shown.
